# PanPPI: ranking-trained protein embeddings for proteome-scale interaction screening

**DOI:** 10.64898/2026.11.27.769337

**Authors:** Brayon J. Fremin, Carlotta Ronda

## Abstract

Proteome-wide interaction screening requires ranking millions of protein pairs before costly structure prediction. PanPPI trains a cacheable protein encoder with a ranking loss, so each protein is embedded once and pairs are scored at 0.75 million per minute on one H200 GPU. PanPPI matched or outperformed published models on benchmarks, and in five eukaryotes its top 1% of pairs contained 52-88% of AlphaFold-Multimer’s confident predictions among STRING-linked pairs.

## Main

Structure predictors such as AlphaFold-Multimer^1^ and AlphaFold 3^2^ can confirm protein-protein interactions (PPIs) but are too costly to run on the 10⁷-10⁸ pairs of a proteome, so cheaper filters come first^3,4^. Sequence-based joint models, which encode each pair together, previously set the accuracy standard^5,6^ but need a forward pass per pair^7^. Bi-encoders embed each protein once and cache the embeddings^7,8^. LoGoPPI^7^, which fine-tunes the language model ESM-2^9^ and adds a late-interaction score (MaxSim)^10^, matches joint models on eukaryotic test sets.

A screen depends on ranking: true partners must score above the many non-partners, as in dense retrieval^11^. Standard benchmarks can also reward memorised hub proteins and training-set similarity^12^ (**Supplementary Fig. 3**), so we tested on the standard benchmark and under leakage-controlled protocols.

PanPPI is a bi-encoder trained with a ranking objective (**Fig. 1a**). One fine-tuned ESM-2 650M encoder feeds two projection heads: proj_g is trained with cross-entropy through a global classifier, and proj_m is trained only with a ranking loss over many non-partners. proj_m is discarded after training; because it is separate from the scoring head, its gradient reaches the score only by shaping the shared encoder. At inference, a global score and a parameter-free MaxSim on proj_g residue embeddings are calibrated and averaged.

**Fig. 1:**
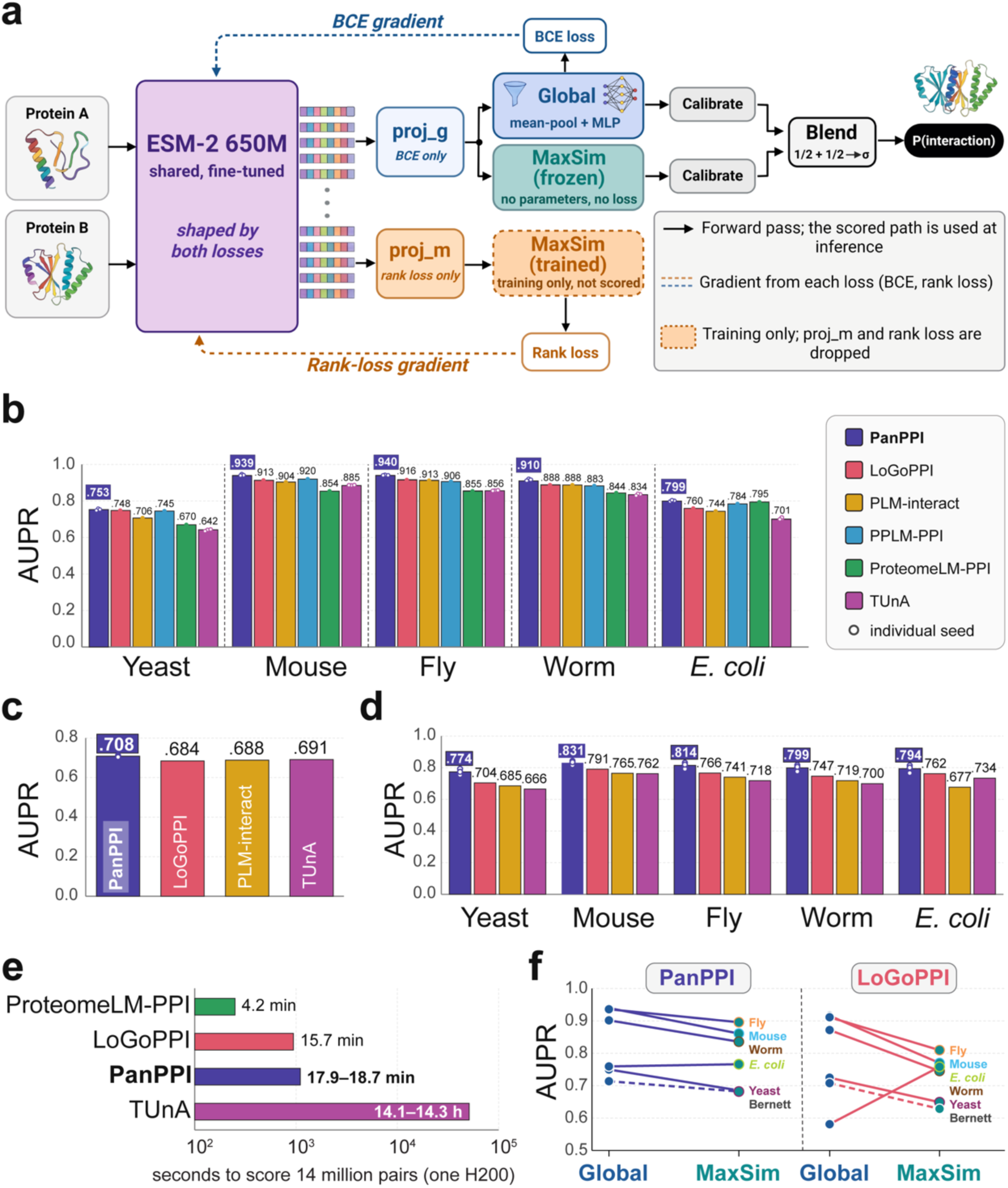
PanPPI design, accuracy and speed. **a**, Architecture: ESM-2 650M encoder feeds proj_g (cross-entropy, global head) and proj_m (ranking loss, training only). **b**, AUPR on the D-SCRIPT cross-species sets, human-trained (per eukaryote, 5,000 positives and about 50,000 negatives; *E. coli*, 2,000 and 16,232). Circles, seeds (PanPPI, TUnA: three); other tools, one released model. **c**, AUPR on the Bernett test set (52,048 pairs, 26,024 positives) for LoGoPPI, PLM-interact, TUnA and PanPPI trained on its training split (PanPPI: three seeds). **d**, AUPR of the Bernett-trained models on cross-species positives with degree-matched negatives (1:1, mean of two replicates; 5,000 positives per eukaryote, *E. coli* 2,000). **e,** Time to score 14 million pairs among 10,000 proteins on one H200. **f**, AUPR of the global and MaxSim branches of PanPPI (left) and released LoGoPPI (right) on the standard sets (solid) and the Bernett test set (dashed).

We first asked whether the ranking objective, the main change from LoGoPPI^7^, improves accuracy. Trained on LoGoPPI’s human pairs and recipe, PanPPI had the highest mean area under the precision-recall curve (AUPR) on all five D-SCRIPT^13^ test species, ahead of LoGoPPI by 0.005-0.040, cutting its error (1 − AUPR) by 17-30% in four species (**Fig. 1b**). Paired bootstrap intervals excluded zero in 22 of 25 comparisons with five published models, including ProteomeLM-PPI^14^; the three that did not were yeast (against LoGoPPI and PPLM-PPI) and *E. coli* (against ProteomeLM-PPI), where PanPPI, averaged over three seeds, was ahead by 0.004-0.008 (**Supplementary Table 1**). The lead held when every model was trained on Bernett’s leakage-controlled split^12^ of HIPPIE interactions (AUPR 0.708 against 0.684-0.691; **Fig. 1c, Supplementary Fig. 6**) and when negatives were matched for protein degree (ahead by 0.036-0.120; **Fig. 1d**).

PanPPI enables proteome-scale use. It scored 14 million pairs among 10,000 proteins in 18-19 min on one H200 GPU, similar to LoGoPPI (15.7 min) and far faster than TUnA (about 14 h; **Fig. 1e**); the gain over LoGoPPI is in accuracy, not speed. A control with the same recipe, data and three seeds but no ranking loss scored 0.028-0.040 AUPR lower in every species, with gains of 0.08-0.12 on the MaxSim branch in eukaryotes (**Fig. 1f; Supplementary Fig. 1**).

We next asked whether PanPPI can shortlist pairs for AlphaFold-Multimer (AFM). The reference was pairs that the AlphaFold Database complexes resource^1,15^ predicts with high confidence among STRING^16^ - linked pairs (interface scores ipSAE ≥0.6 and pDockQ2 ≥0.23; ‘AFM-confident’), ranked among all pairs of each proteome (**Fig. 2a**). Because AFM is itself a prediction, this measures agreement with AFM, not experimental truth. In human (pairs new to the model, see **Methods**), yeast, mouse, fly and worm, PanPPI’s top 1% held 52-88% of AFM-confident pairs (LoGoPPI 43-83%; paired differences 3-14 points, intervals excluding zero), against 1% expected by chance (**Fig. 2b**). With the 10% most-connected proteins removed, PanPPI still held 51-83% (**Supplementary Table 14, Supplementary Fig. 8**). At a fixed budget of 10⁴ pairs sent to AFM, PanPPI found 2.0-4.6 times as many AFM-confident pairs as LoGoPPI in these five species (103 against 26 in human), and 1.1-1.3 times as many at 10⁶ (**Supplementary Table 18**). Pairs with weak STRING support (scores below 400), 35% of the AFM-confident pairs in human, were retained at 35-80%, 6-18 points above LoGoPPI (**Supplementary Fig. 7**). In *E. coli,* retention was weaker for both tools (PanPPI: 14% and LoGoPPI: 19%; **Supplementary Note 1**). Five further species (soybean, *Arabidopsis*, rat, rice, zebrafish) gave 73-78% in the top 1%, 4-7 points above LoGoPPI (**Supplementary Fig. 9**).

**Fig. 2:**
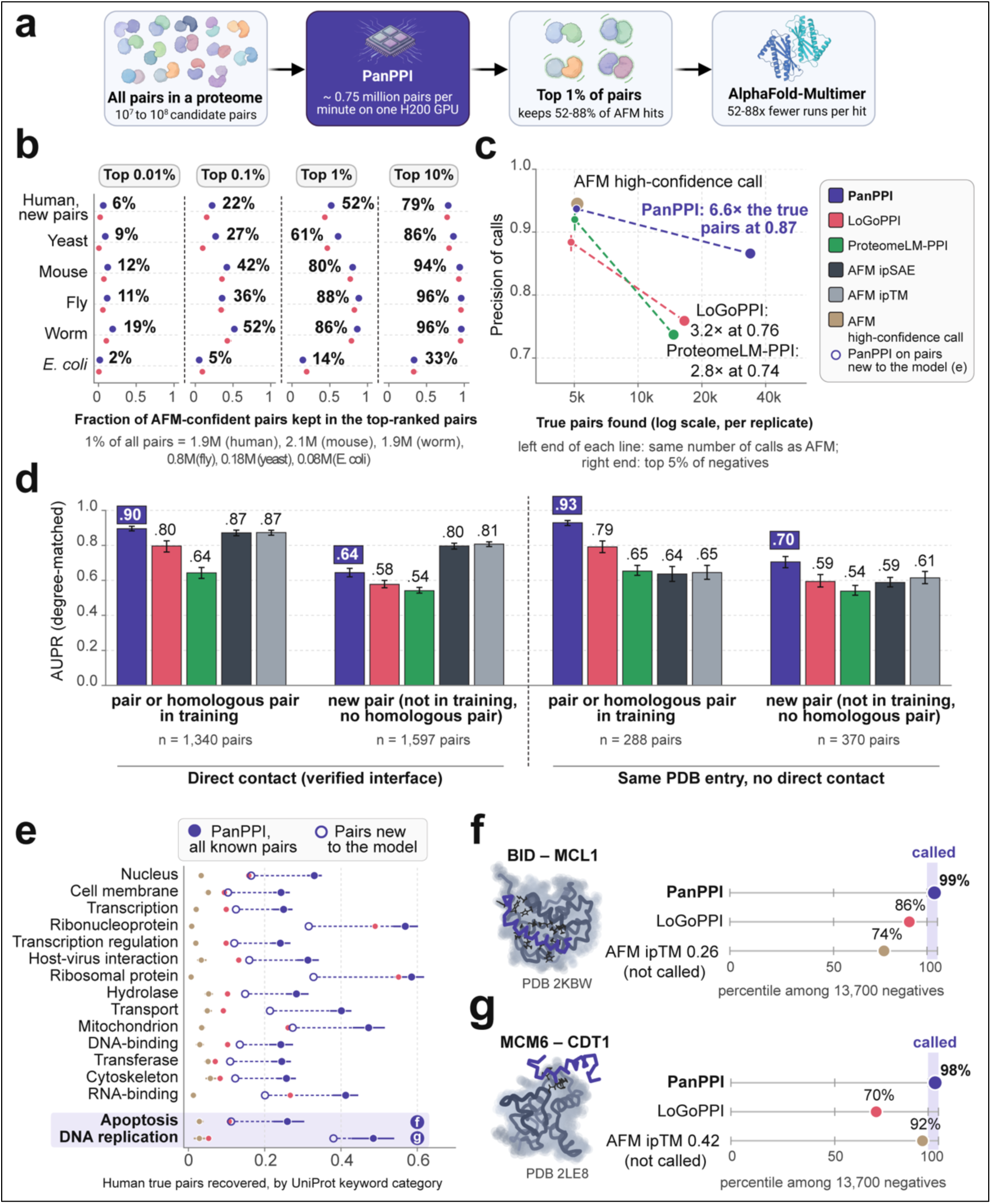
PanPPI as a first stage before AlphaFold-Multimer (AFM). **a,** Workflow. **b,** AFM-confident pairs (ipSAE ≥0.6 and pDockQ2 ≥0.23) kept in the top 0.01%, 0.1%, 1% and 10% of all pairs (human: pairs new to the model); 95% bootstrap intervals; numbers of pairs in Supplementary Table 18. **c,** Precision against STRING physical (≥700) pairs versus true pairs found; left ends, as many calls as the AFM high-confidence call; right ends, above the 95th percentile of each method’s negatives; nine evaluation sets pooled, three negative replicates. **d,** Degree-matched AUPR for human pairs in one PDB entry, with or without direct contact, by training overlap (with contact: 1,340 seen, 1,597 new; without: 288 seen, 370 new; 15,852 negatives). **e,** Known human pairs recovered by UniProt category (open: new to the model). **f, g,** Two examples of contacts called by PanPPI but not AFM, selected as in Methods (ipTM 0.26, 0.42; ipSAE 0.00, 0.04); percentile among negatives.

The ranking surfaced pairs that AFM’s confident calls miss. When PanPPI called as many pairs as AFM, STRING precision was similar (0.937 against 0.945); calling every pair above the 95th percentile of its negatives found 6.6 times as many STRING pairs, at lower precision (0.866 against AFM’s 0.945; **Fig. 2c,e**). STRING-derived labels may favour sequence models trained on related data and, outside human, may be partly transferred from human by orthology; AFM is neither. On experimental structures, PanPPI ranked human pairs sharing a PDB entry above degree-matched non-pairs, including new ones (AUPR 0.64 with direct contact, 0.71 without; chance 0.50, LoGoPPI 0.58 and 0.59). AFM was better for direct contacts (0.80), PanPPI for same-complex pairs without contact (0.59 for AFM; **Fig. 2d,f,g**).

PanPPI is a fast first stage that keeps about half to most of what AFM would confirm in eukaryotes and finds further STRING-supported pairs, while AFM remains the better judge of direct contact. The evidence is retrospective: AFM was run only on STRING-linked candidates, precision is measured on balanced sets rather than at proteome prevalence, and finding interactions with no prior evidence was not tested, so PanPPI is best used to prioritize pairs for confirmation by AFM or experiment. Contrastive retrieval models built for microbial proteomes^17^ or structure-based inputs^18^ were not benchmarked (**Methods**). Choose the top fraction by AFM budget (**Fig. 2b**); the gain over LoGoPPI is largest when that budget is small (**Supplementary Table 18**). PanPPI is released as a command-line tool, panppi-score, with all trained models (**Code availability**).

## Methods

### Terms

*AFM-confident* pairs carry the high-confidence label of the AlphaFold Database complexes resource (ipSAE ≥ 0.6 and pDockQ2 ≥ 0.23). A pair is *new* to PanPPI if neither the pair nor a training positive pair linking homologs of both proteins was in its training set (strata 3-5); *no homolog* means no human training protein at ≥ 40% identity and ≥ 50% query coverage. A *hub* is a protein in the top share of the distribution of STRING-linked AlphaFold Database candidate partners per protein. *Retention* at x% is the share of AFM-confident pairs ranked in the top x% of all proteome pairs. *Degree-matched* negatives contain each protein as often as the positives do.

### Architecture

PanPPI (Partner-ranking Alignment Network) is built on an ESM-2 650M^9^ encoder (hidden size 1,280), fine-tuned end to end on the first 800 residues of each protein; the contact head and pooler are frozen because they do not reach the loss. For a protein A the encoder returns residue embeddings H_A_ ∈ ℝ^|A|×1280^ (start and end tokens removed). Two linear layers map them to 512 dimensions, each followed by dropout (0.1): proj_g gives X_A_ = Drop(W_g_H_A_) and proj_m gives Y_A_ = Drop(W_m_H_A_), with weights initialised from N(0, 0.02). Scoring uses X only.

The global branch mean-pools X over residues, p_A_ = |A|^−1^ Σ_i_ X_A,i_, and forms g_AB_ = LN([p_A_ ǁ p_B_ ǁ |p_A_ − p_B_|]) ∈ ℝ^1536^, where ǁ is concatenation and LN is layer normalisation. A two-layer classifier (1,536 → 512, ReLU, dropout 0.1, 512 → 1) gives the logit ℓ_AB_, and the symmetrised global logit is

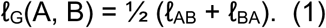

The local branch is the parameter-free symmetric MaxSim^10^ on the same embeddings,

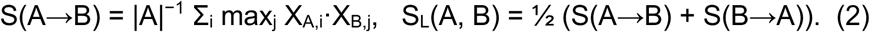

Both branches are symmetric in A and B by construction, so a pair’s score does not depend on the order of its proteins.

### Ranking objective

proj_m is used only in training. Within each per-device micro-batch of pairs (A_i_, B_i_), i = 1…n (n = 16-24), with labels y_i_, compute s_ij_ = S_L_(Y_Ai_, Y_Bj_) between every first protein and every second protein. For each positive pair (y_i_ = 1) the true partner must outscore the other proteins in the batch, with known training positives of A_i_ masked out:

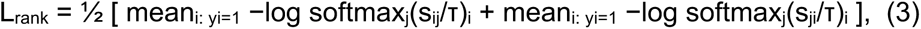

with temperature τ = 1 and raw (unscaled) MaxSim scores. The total loss is L = L_BCE_ + λ L_rank_, where L_BCE_ is binary cross-entropy on ℓ_G_ (positive-class weight 10 for the human data and 1 for Bernett) and λ = 1. The gradient of L_BCE_ reaches proj_g and the encoder, and that of L_rank_ reaches proj_m and the encoder, never proj_g; proj_m is discarded after training. The control model uses the same code, data and selection rule with λ = 0, which is the LoGoPPI recipe^7^.

### Calibration and scoring

After training, the encoder and heads are frozen and each branch is calibrated on a held-out human calibration split. The global logit is rescaled as ℓ̃_G_ = ℓ_G_/T + b_G_, and MaxSim, after standardisation with its mean μ and standard deviation σ over the training pairs, as ℓ̃_L_ = a (S_L_ − μ)/σ + b_L_, with a, T > 0. The four parameters are fitted by minimising the unweighted negative log-likelihood (L-BFGS-B, log T and log a bounded to [−10, 10]). The final logit is the equal-weight average

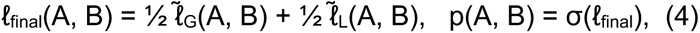

with no fitted blend weight. Each protein is embedded once, X is cached, and pairs are scored from the cache.

### Training

Human training pairs, validation splits and recipe follow LoGoPPI^7^, which built its human set from 47,932 positive pairs (STRING v11 physical binding interactions with experimental evidence, following D-SCRIPT^13^) split 80/20 into training and validation, with negatives drawn at random at 10:1 (**Supplementary Tables 8, 11 and 12**). Our training file has 421,312 pairs (38,267 positive; LoGoPPI reports 38,345 before removing duplicate and conflicting pairs). We used AdamW with a learning rate of 2×10⁻⁵, 10% warmup and linear decay, weight decay 0.01, gradient clipping at 1.0, bfloat16, and data-parallel training with gradient accumulation to an effective batch size of 96 in every run (two GPUs × 24 × 2 for seeds 42 and 7; two GPUs × 16 × 3 for seed 141; one GPU × 24 × 4 for the Bernett runs). The positive-class weight was 10 for the 1:10 human data and 1 for the 1:1 Bernett data. Training ran for at most 14 epochs with length-grouped batches. After each epoch the calibrated blend was scored on a selection split (a held-out part of the human validation pairs, or half of the Bernett validation pairs), the best epoch was kept, and training stopped after three epochs without improvement. Every run stopped this way and none reached the 14-epoch ceiling: PanPPI trained on human data for 9, 8 and 11 epochs (best epochs 6, 5 and 8; seeds 42, 7, 141), the control for 11, 11 and 8 epochs (best 8, 8 and 5), and PanPPI on the Bernett split for 6, 5 and 8 epochs (best 3, 2 and 5; **Supplementary Fig. 5 and Supplementary Table 8**). Seeds were 42, 7 and 141.

### Uncertainty and familiarity layers

Three label-free layers were added after training and are optional (**Supplementary Note 2, Supplementary Fig. 4, Supplementary Tables 6 and 9**). (i) Pair-level GP variance: a random-feature Gaussian process with 1,024 Fourier features (amplitude set by the median heuristic) is fitted to the penultimate features of the global classifier on the training pairs, with the precision matrix accumulated without labels (leverage form, ridge penalty 10⁻³), and its predictive variance for a pair is the model-uncertainty feature. (ii) Protein-level familiarity: for each protein, the mean cosine distance between its pooled projected embedding and those of its ten nearest training proteins; a pair contributes the smaller and the larger of its two distances. (iii) MaxSim evidence: the mean gap between each residue’s best and second-best match in the partner, the spread of the best-match scores, and the asymmetry of both between the two directions. B0 is the calibrated blend z; B1 adds log GP variance; B2 adds the two distances; B3 adds the MaxSim statistics. Each is a logistic regression on standardised features (L2 penalty, C = 1), fitted on held-out sets as stated in the text of each analysis and evaluated on pairs the regression has not seen, with the same paired, protein-clustered bootstrap as above. To ask whether the GP variance flags errors, an error was defined as misclassification of the calibrated blend at probability 0.5, and for each set and seed we computed the AUROC of the variance, and of low confidence (−|logit|), for predicting errors, with the Pearson correlation between variance and |logit|.

### Metrics

AUPR is the average precision with the interaction class positive; sets are evaluated at 1:1 prevalence unless stated. Retention at rank fraction x is the share of AFM-confident pairs whose score is within the top fraction x of all proteome pairs. For degree-matched evaluation of verified contacts, each pair is given d = log(1 + k_A_) + log(1 + k_B_), where k is the protein’s degree in the evaluated set; negatives in bin b of d (20 bins from the quantiles of the positives) receive weight w_b_ = π^+^_b_/π^−^_b_, where π is the share of each class in the bin, rescaled so that the weights sum to the number of positives. The degree-only AUROC of the weighted set is 0.50 by construction. A tool’s error is 1 − AUPR, and the relative error reduction is 1 minus the ratio of PanPPI’s error to the comparator’s.

### Statistics

PanPPI results are the mean of three training seeds, with individual seeds shown, capturing training-seed variation. Test-set uncertainty was estimated with a paired, protein-clustered bootstrap (1,000 resamples): each pair was assigned to its higher-degree protein, proteins were resampled with replacement, and every tool was scored on the same resample. AUPR was averaged over the three seeds of PanPPI, and over the three released seeds of TUnA in panel b, within each resample. Intervals are the 2.5th and 97.5th percentiles of the paired difference (PanPPI minus tool) and cover test-set noise only; tools other than PanPPI and TUnA are single models, so their seed variation is not captured. Panel d intervals use the first of the two negative replicates; panel c uses the 51,978 of 52,048 test pairs scored by all tools. Full results are in **Supplementary Table 1. For Fig. 2**, retention intervals use 300 resamples of proteins (each AFM-confident pair assigned to its higher-degree protein), the PanPPI minus LoGoPPI difference is taken on the same resample, the supplementary retention analyses (hub removal, STRING tiers) use 300 resamples, and call precision and contact AUPR use 300 resamples within each set with all tools scored on the same resample.

### Standard cross-species benchmark

Human-trained models were tested on the D-SCRIPT test sets for yeast, mouse, fly, worm and *E. coli*^13^, in the curated PPLM-PPI versions that LoGoPPI uses^7^ (5,000 positives and 49,963-49,999 negatives per eukaryote; *E. coli* 2,000 positives and 16,232 negatives, prevalence 0.110). Metrics are AUPR and AUROC on the full test files. AUPR depends on prevalence, so values are comparable only on identical files.

### Comparators

We scored all tools on identical pair files using released models (**Supplementary Table 17**): LoGoPPI-Cross-species with the authors’ inference script, PPLM-PPI and PLM-interact with their released weights, ProteomeLM-PPI with its supervised score^14^, and TUnA with its released checkpoints for seeds 47, 94, and 141. Where published values exist our scores reproduce them (for example PPLM-PPI AUPR 0.745 in yeast and 0.784 in *E. coli*). For the Bernett comparisons, we used the released Bernett-trained checkpoints of LoGoPPI, PLM-interact and TUnA; PPLM-PPI and ProteomeLM-PPI are not shown because no Bernett-trained checkpoints exist. Two contrastive retrieval models trained on structure-derived contact labels were not benchmarked because their training data and tasks differ from the interaction-pair setting evaluated here.

FlashPPI^17^ is a dense-retrieval model with contact-map reranking, built on a genomic language model pretrained mainly on prokaryotic metagenomes; it is trained on interface contacts from PDB complexes and AlphaFold Database domain–domain interactions and was evaluated on *E. coli* and prokaryotic host–virus systems. ColBERT-PPI^18^ is a late-interaction model that takes a predicted structure for every protein (Foldseek 3Di tokens) and is trained on residue contacts from experimental complexes (PINDER). PanPPI and the five comparators instead learn from interaction-pair labels and are evaluated here on interaction-pair benchmarks. Scoring the released retrieval models on these sets would test them on a label type, and for FlashPPI on taxa, outside their design, and a like-for-like comparison would require retraining all models on common data. PanPPI needs sequence only, with no structures or alignments in training or inference, applies its ranking loss to an auxiliary head that is discarded after training, and was evaluated in ten eukaryotic proteomes from animals, plants and yeast. The released LoGoPPI is a single run (seed 42)^7^, so its seed-to-seed variation is unknown; PanPPI’s three-seed mean exceeded it in all five species (**Fig. 1b**).

### Bernett benchmark

We used the training, validation and test splits of Bernett et al.^12^ (test: 52,048 pairs, 26,024 positives). PanPPI was trained on the Bernett training split with the recipe above (seeds 42, 7, 141).

### Degree-matched cross-species sets

For each species the original test positives were kept. Negatives were constructed so that every protein appears in the negatives exactly as often as in the positives, by a Havel-Hakimi construction with degree-preserving swaps, excluding positive pairs, duplicates and self-pairs; two replicates were built. The degree-only AUROC of these sets is 0.50-0.53.

### Hard-negative evaluation

Following LoGoPPI’s design, a negative pair is hard if its two proteins share a direct GO cellular-component annotation (GOA human, aspect C; excluding the root, terms of the subset gocheck_do_not_annotate, and protein-containing complex with its is_a descendants; go-basic version 2026-07-26) or an identical Pfam accession (UniProt release 2026_04; **Supplementary Note 4, Supplementary Fig. 6, Supplementary Table 10**). We evaluated the Bernett test set with Bernett-trained models. The positives are drawn with replacement from the positives that also share the annotation, up to the number of hard negatives; hard negatives and size-matched random negatives are resampled with replacement (2,000 replicates, the same positives in both comparisons); PanPPI is the mean of its three seeds’ AUPR.

### Speed benchmark

All tools were run on 14 million pairs among 10,000 proteins sampled from mouse, on one H200 GPU, with protein representations computed once and reused. Times include protein embedding and pair scoring, and model loading for LoGoPPI and PanPPI (ProteomeLM-PPI: embedding and scoring). LoGoPPI, PanPPI, ProteomeLM-PPI, and TUnA were timed on the full 14 million pairs. The released PanPPI tool also has an all-vs-all mode that keeps all embeddings on the GPU and scores one protein against many partners per matrix multiplication; it returns the same scores (differences below 10⁻⁵ logits in its self-check) and scored about 31,000-34,000 pairs per second on one RTX A5000 or A40 (scoring step only). Hence, the benchmark rate is conservative for proteome-scale use.

### Software and hardware

PanPPI was implemented in Python 3.12.8 with PyTorch 2.9.0 (CUDA 12.8) and Hugging Face transformers 5.22.2. LoGoPPI was run in a separate environment with transformers 4.57.6, because its model class relies on a weight-tying attribute renamed in transformers 5, and PLM-interact with transformers 4.40.1 and sentence-transformers 2.7.0. Training used bfloat16 autocast on NVIDIA H200 GPUs (two per job for the human-trained runs, one for the Bernett runs), with an effective batch of 96 in every run; the benchmark files and the speed benchmark were scored with the encoder in bfloat16 autocast and the head in 32-bit floats, and the degree-matched cross-species sets and the proteome-scale scores with 32-bit embeddings (the two agree to a mean of about 0.01 logits and the same AUPR). The speed benchmark used one H200.

### AlphaFold-Multimer reference

AFM scores (ipTM, ipSAE, pDockQ2) and the high-confidence call come from the AlphaFold Database complexes resource^15^ (AlphaFold-Multimer model_1_multimer_v3, four recycles with early stopping, no relaxation; monomer alignments concatenated without pairing) for STRING^16^ physical pairs without a score threshold. AFM-confident pairs are those carrying the resource’s high-confidence label, defined as ipSAE ≥ 0.6 and pDockQ2 ≥ 0.23, each taken as the maximum over the two directions^15^; the authors report a precision of 0.958 and a false-positive rate of 0.004 for it on a PDB heterodimer benchmark (94 positives, 250 controls). In the metadata release we used (17 July 2026), 80,248 of 7,594,357 heterodimer candidates carry the label, and the rule reproduces it on every row.

### Degree-balanced proteome-scale sets

Positives were STRING v12 physical interactions with a combined score of at least 700 among the proteins of each AlphaFold Database proteome (**Supplementary Fig. 2**). Negatives were protein pairs absent from the STRING physical network, built so that each protein occurs in the negatives as often as in the positives (Havel-Hakimi construction with degree-preserving swaps; three replicates), which removes the protein-frequency shortcut that inflates pair-level benchmarks. Residual degree signal is reported as the AUROC of a degree-only score (0.52-0.60 for all proteins, 0.51-0.58 for proteins without a training homolog; “no homolog” = no human training protein at 40% identity and 50% coverage by MMseqs2).

### Sets for the comparison with AlphaFold-Multimer

For these comparisons, negatives were drawn from low-confidence AlphaFold Database candidates (STRING below 400, no PDB co-membership), which constrains matching; positives were trimmed to those whose degrees the candidate graph could match (38-72% of the STRING positives were retained in the sets used), AFM-confident pairs were not removed from the negatives (excluding them changed AUPR by at most 0.02), and sets were used only if the negatives reached 80% of the retained positives and the degree-only AUROC was at most 0.60 (**Fig. 2c**); Supplementary Table 15 gives the comparison without trimming.

### Retention, call precision and categories

Retention is the fraction of AFM-confident pairs whose rank among all pairs of the proteome is within the top x%, averaged over PanPPI seeds; the pairs are all unordered pairs of the proteins in each species file (human 19,408; mouse 20,612; fly 12,786; worm 19,734; yeast 5,960; *E. coli* 4,091 proteins), and no degree control is applied to retention (**Supplementary Tables 2, 3 and 16**; a hub-trimmed version, which removes pairs involving proteins in the top 1%, 5% or 10% of candidate-partner degree, is in **Supplementary Table 14**; **Supplementary Fig. 8** repeats the retention analysis with ipSAE or ipTM alone as the definition, for each seed separately, after hub removal and by training novelty, with 200 protein-clustered resamples for the intervals; the five further species, built from the same public resources and evaluated with the same definitions, are described with **Supplementary Fig. 9**). Call precision pools the sets above (one species at one homology level) and three negative replicates; matched-count calls take, per set, the top k pairs of each tool, where k is the number of AFM calls, and threshold calls take every pair above the 95th percentile of the tool’s own negatives in that set. The percentile among negatives in **Fig. 2f,g** is the share of the 13,700 negatives scored below the pair. Keyword categories are the 14 largest non-disease UniProt keywords of either protein, and recall uses each tool’s default operating point. Training-overlap labels use MMseqs2 (sensitivity 7.5, E ≤ 10⁻³) hits at 40% identity and 50% query coverage, with exact matches at ≥99.9% identity and ≥99% coverage. The two examples are illustrations chosen from direct contacts new to PanPPI that AFM did not call; they are not a random sample.

### Fixed-budget counts

For a budget of B pairs sent to AFM (10⁴, 10⁵ and 10⁶), the number of AFM-confident pairs found was the number whose blend rank among all pairs of the proteome was at most B divided by the number of pairs of that proteome; PanPPI is the mean over its three seeds and LoGoPPI is the released model. Intervals are protein-clustered bootstrap percentiles of the difference (200 resamples, each pair assigned to its higher-degree protein), as for retention. Only pairs with an AFM result are counted, so the numbers are lower bounds for both tools (**Supplementary Table 18**).

### Cost per AFM-confident hit

For a species with N proteins in the screened file the universe is N(N − 1)/2 pairs (**Supplementary Table 2**). A top fraction x sends x of the universe to AFM, retention r is measured as above, and the cost per hit is x × universe/(r × n), where n is the number of AFM-confident pairs of the set; random selection costs universe/n. AFM-confident pairs are known only among STRING-linked candidates, so n is a lower bound on the true number and the cost per hit an upper bound. Retention at 0.01%, 0.1%, 1% and 10% was averaged over PanPPI seeds, and intervals use 500 protein-clustered resamples with PanPPI and LoGoPPI scored on the same resample.

### Homology searches and training overlap

Proteins were compared with MMseqs2 (release 15-6f452; easy-search, sensitivity 7.5, E ≤ 10⁻³; up to 500 hits per query for the search against the human training proteins and 2,000 for the search against PDB chains; Supplementary Fig. 3, Supplementary Table 5). Two proteins were called homologous at ≥40% identity and ≥50% query coverage, and identical at ≥99.9% identity and ≥99% coverage of both sequences. Species-level “no homolog” sets keep only pairs in which neither protein has a homolog among the human training proteins. For human pairs, stratum 1 is a pair present in the training set (as a positive or a negative), stratum 2 a pair that is absent but linked to a training positive pair through homologs of both proteins, and strata 3-5 pairs with neither; strata 3-5 are called new, and may still contain proteins with homologs elsewhere in training.

### PDB co-membership and contacts

PDB-tier positives are pairs of human proteins found in one PDB entry, with chains mapped to UniProt accessions through SIFTS. For entries with at most 8 chains, coordinates were taken from the RCSB PDB, and a pair has a direct contact in an entry when at least three residues of each protein have an atom within 4.5 Å of an atom of the partner (KD-tree neighbour search). A pair is verified if it has a direct contact in any such entry; the others are co-members without contact.

### Verified contacts

Verified contacts were PDB co-memberships (entries of at most 8 chains) with at least three residues of each protein within 4.5 Å; negatives were reweighted so their degree distribution matched the positives’ (degree-only AUROC 0.50), and AUPR is reported at equal numbers of positives and negatives (**Fig. 2d, Supplementary Table 4**). Pairs were labelled as seen by PanPPI if the pair itself, or a training positive pair linking homologs of both proteins, was in its training set, and as new otherwise. Intervals are paired, protein-clustered bootstrap percentiles as above. Pairs that share a protein are not independent, so each pair is assigned to its higher-degree protein as the resampling unit; this approximation ignores dependence through the second protein.

### Bernett-trained models on verified contacts

The Bernett-trained PanPPI and LoGoPPI were scored on the same 30,024 human candidate pairs, and the same positives, negatives and strata, as **Fig. 2d**, with AUPR computed as for that panel and a protein-clustered bootstrap of 300 resamples in which all tools were scored on the same resample (**Supplementary Note 3, Supplementary Table 7**).

### Keyword categories

Each protein was annotated with its UniProt keywords, and a pair belongs to a category if either protein carries the keyword. Categories were keywords with at least 800 STRING-positive human pairs after removing generic annotation terms (for example *Reference proteome*, *Phosphoprotein*, *Membrane*) and disease keywords; the 14 largest are shown, plus the categories of the two examples. Recall is the share of the pairs in a category that a tool calls at its default operating point (the AFM high-confidence call, or a score above the 95th percentile of each replicate’s negatives for PanPPI and LoGoPPI, averaged over replicates). Intervals are 95% percentiles of 300 bootstrap resamples clustered on the first protein of each pair.

### Selection of examples

The two examples (PDB 2KBW and 2LE8) were chosen from verified contacts with an interface of at least 10 residues per protein, entries of at most four chains and a STRING physical score of at least 700, for which AFM gave no high-confidence call and an ipTM below 0.5, PanPPI scored above the 95th percentile of the negatives in at least two of three replicates and LoGoPPI in at most one, favouring well-characterised pairs that are new to PanPPI. All pairs meeting these criteria are listed in Supplementary Table 13. They illustrate the behaviour and are not a random sample.

## Data availability

STRING v12, Gene Ontology annotations and UniProt cross-references, the D-SCRIPT cross-species sets, the Bernett splits, the AlphaFold Database complexes resource and the RCSB PDB are public; trained weights, derived per-pair scores and annotation tables will be deposited on Zenodo.

## Code availability

The ‘panppi-score’ tool and training and analysis code will be made available. Trained weights will be deposited on Zenodo.

## Author contributions

Conceptualization and discussion: B.J.F., C.R. Model development, data curation and analysis: B.J.F. Resources: C.R. Writing, review and editing: B.J.F. and C.R.

## Acknowledgements

C.R. acknowledges funding support in part from Lyda Hill Philanthropies, Acton Family Giving, the Valhalla Foundation, Hastings/Quillin Fund - an advised fund of the Silicon Valley Community Foundation, the CH Foundation, Laura and Gary Lauder and Family, the Sea Grape Foundation, the Emerson Collective, Mike Schroepfer and Erin Hoffman Family Fund - an advised fund of Silicon Valley Community Foundation, the Anne Wojcicki Foundation through The Audacious Project at the Innovative Genomics Institute; the Food Allergy Fund and Sandia National Lab.

## Additional information

Correspondence and requests for materials should be addressed to Carlotta Ronda. Supplementary information accompanies this paper.

---

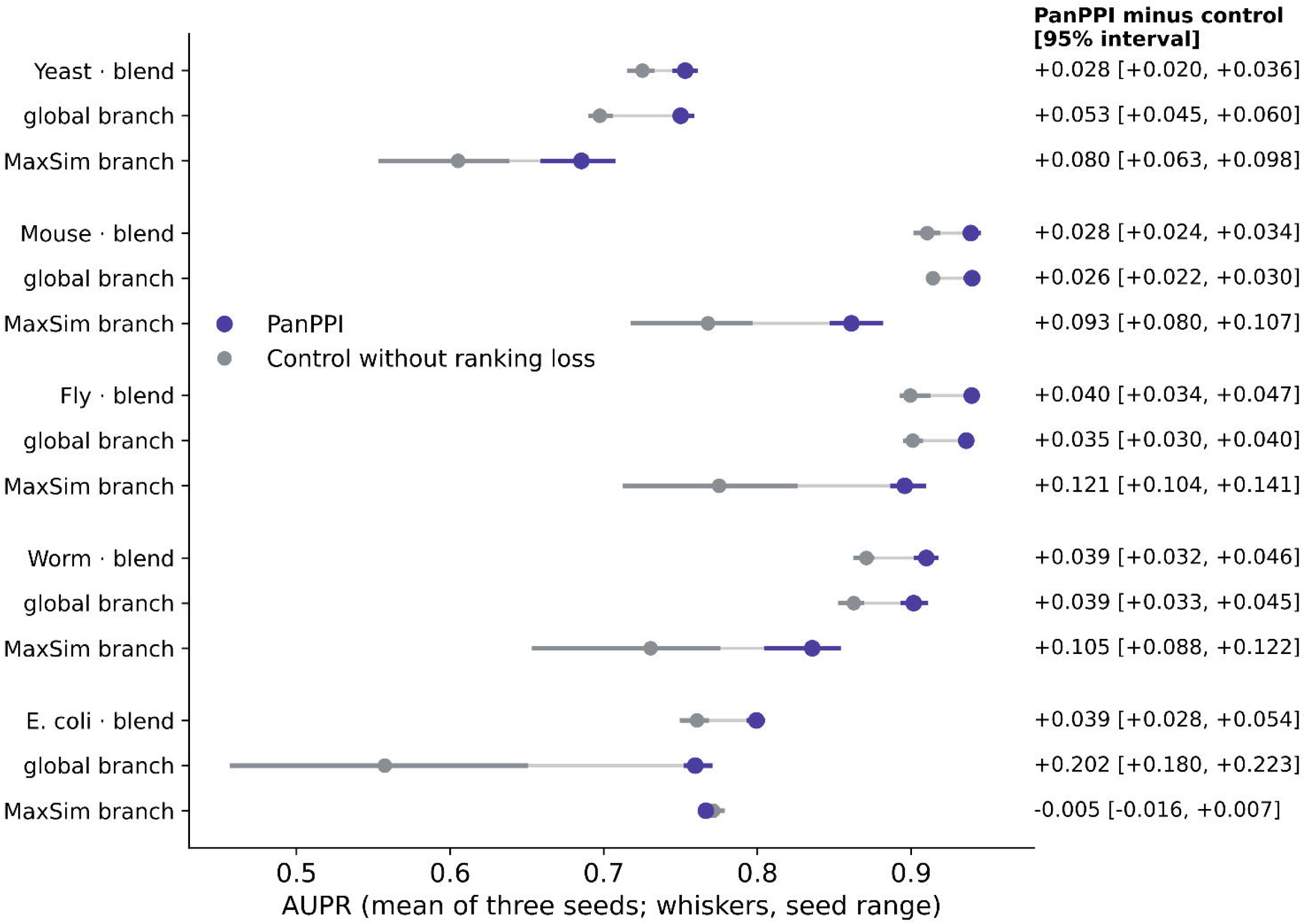

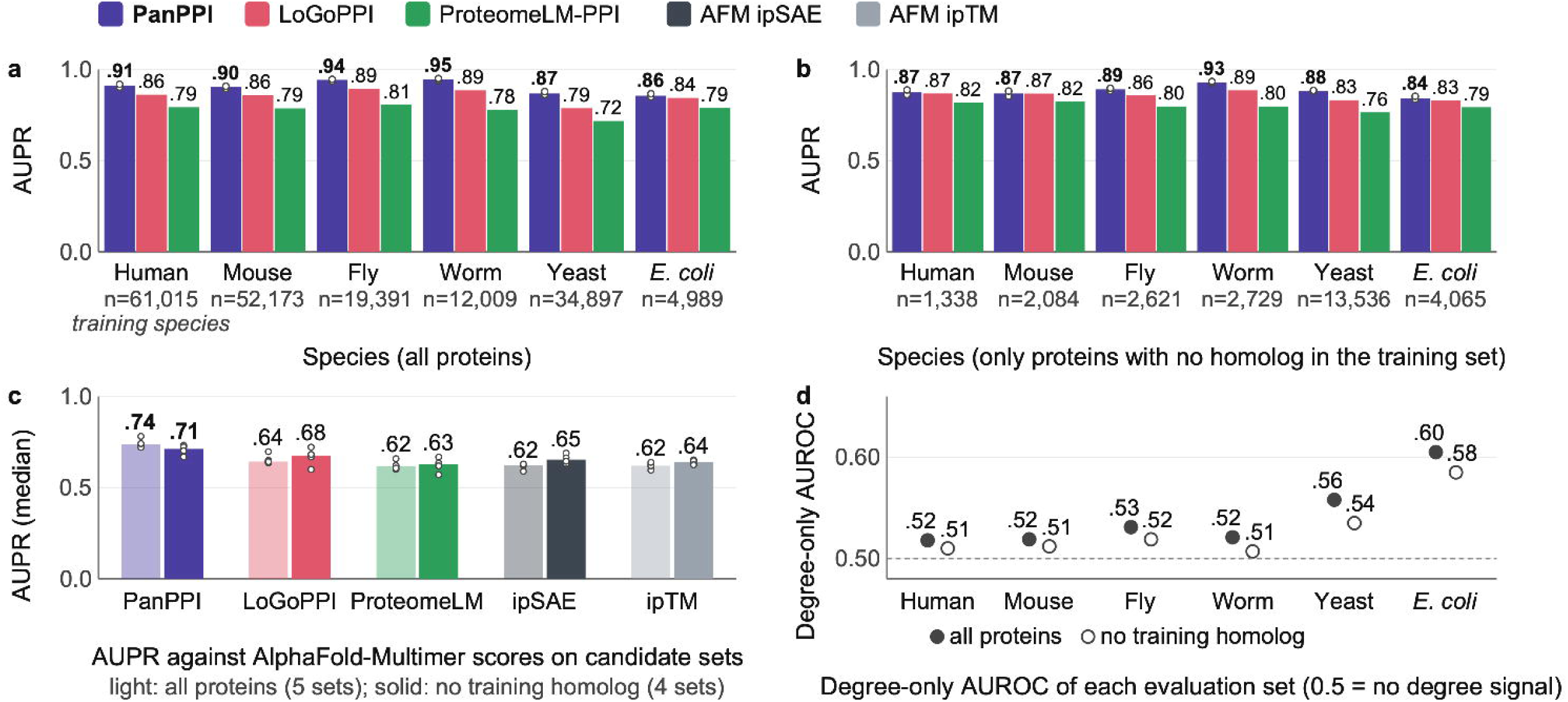

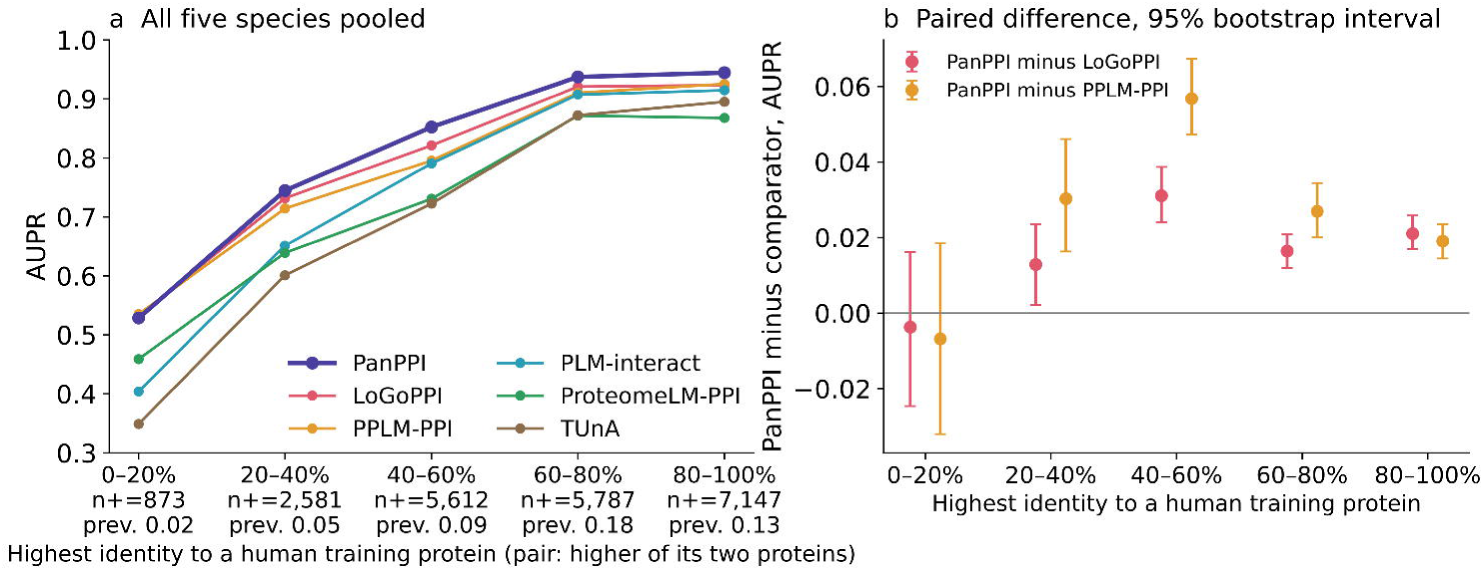

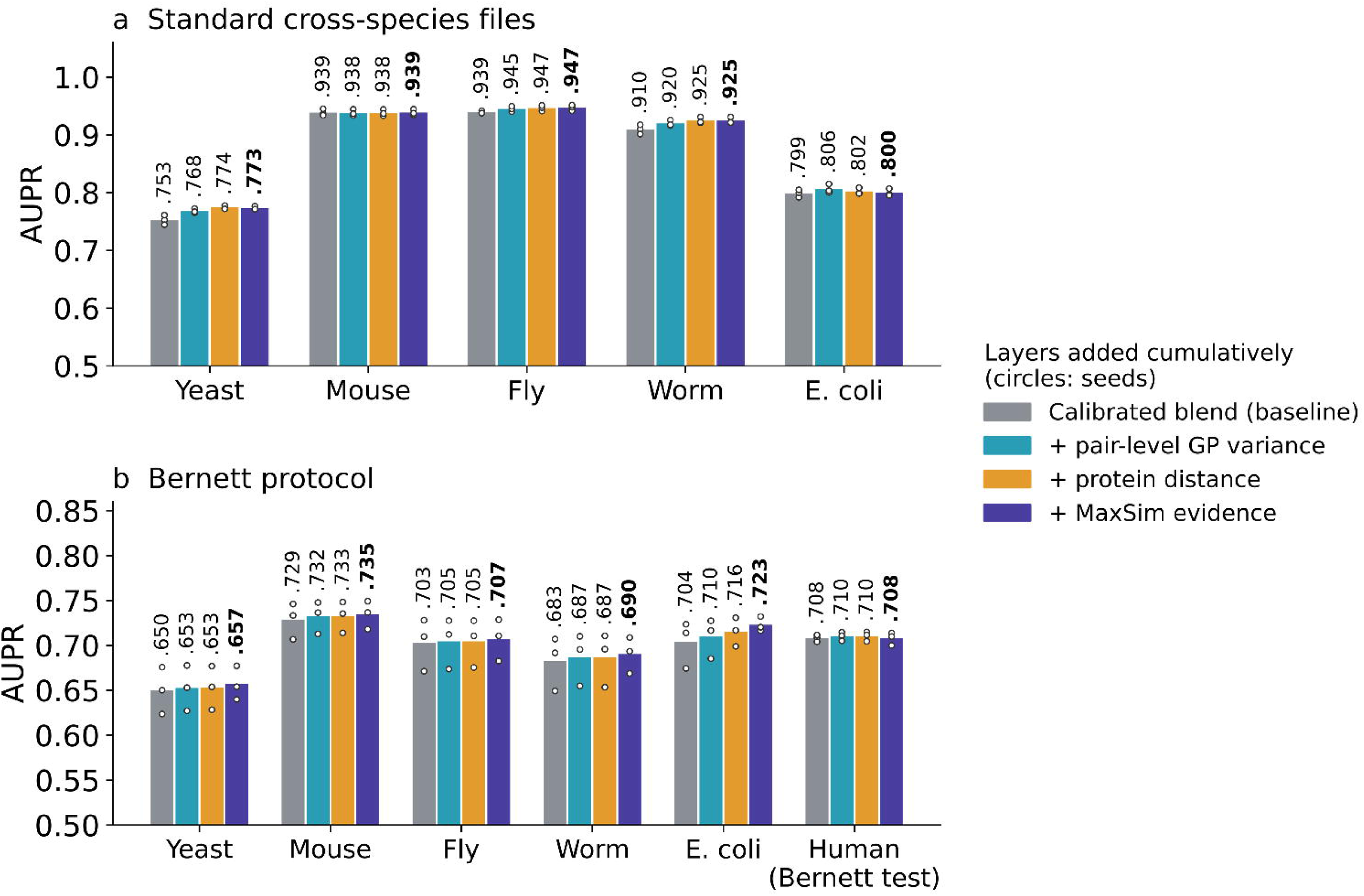

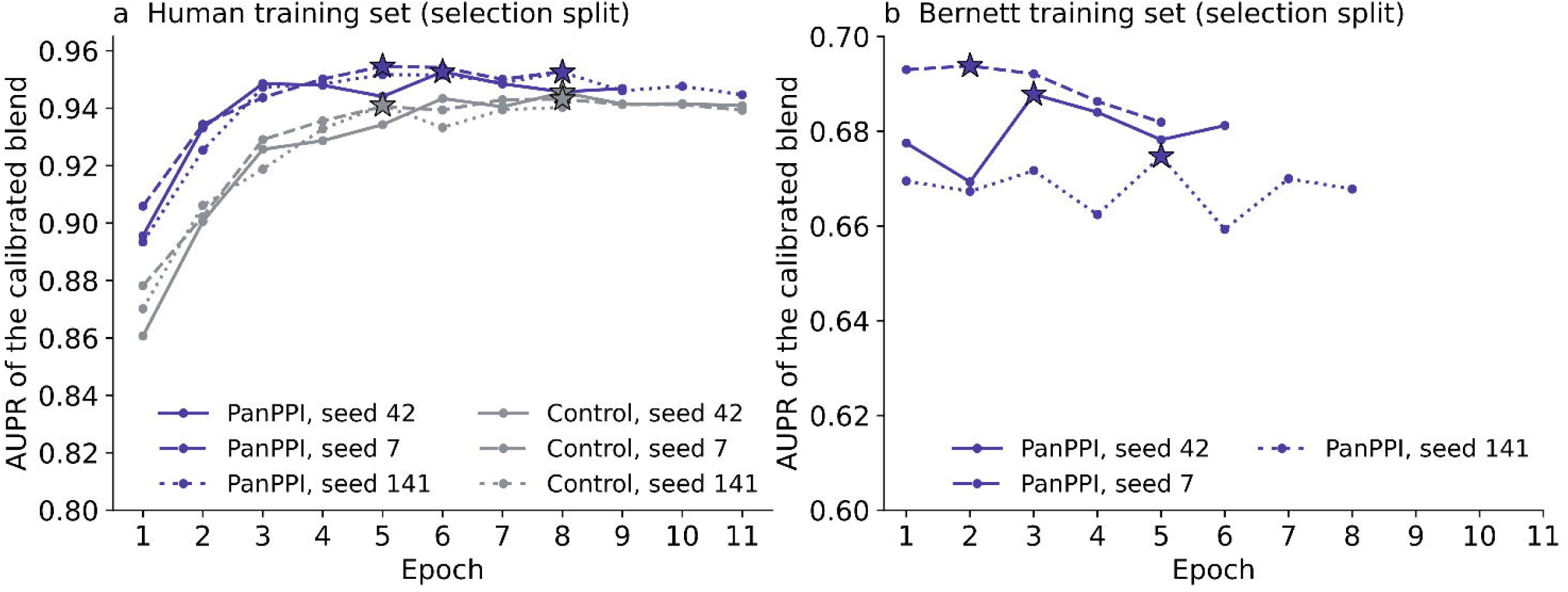

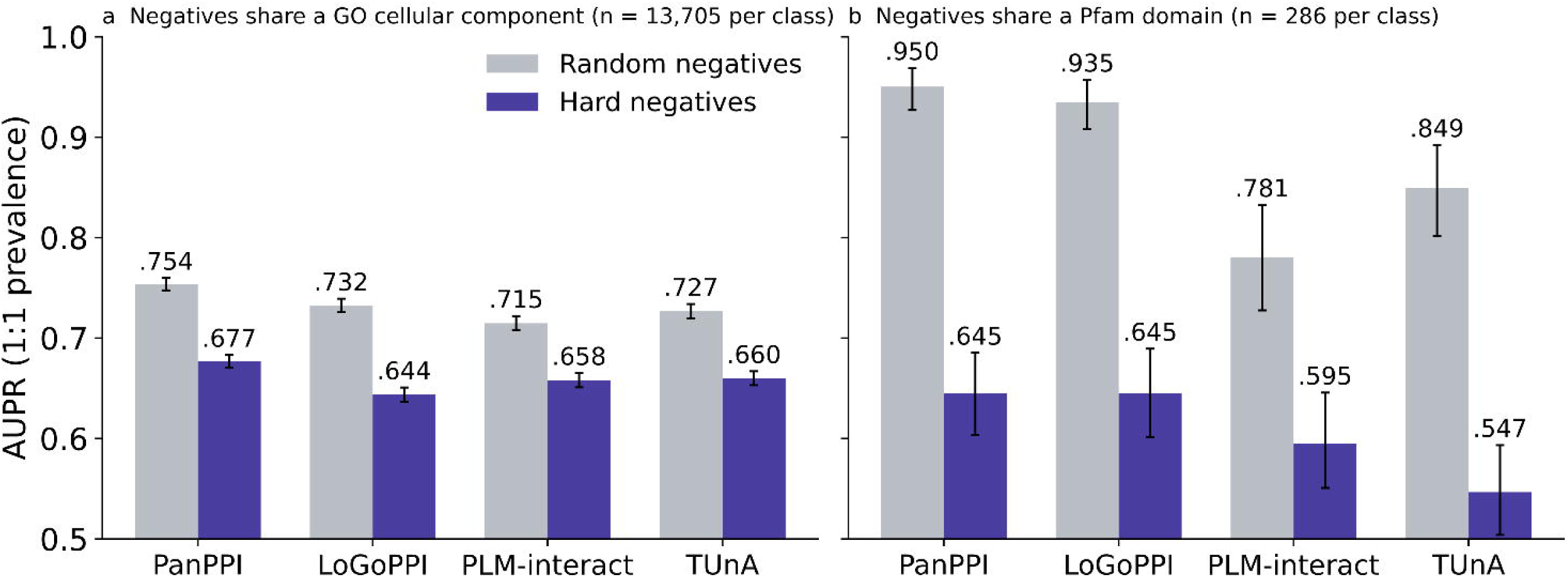

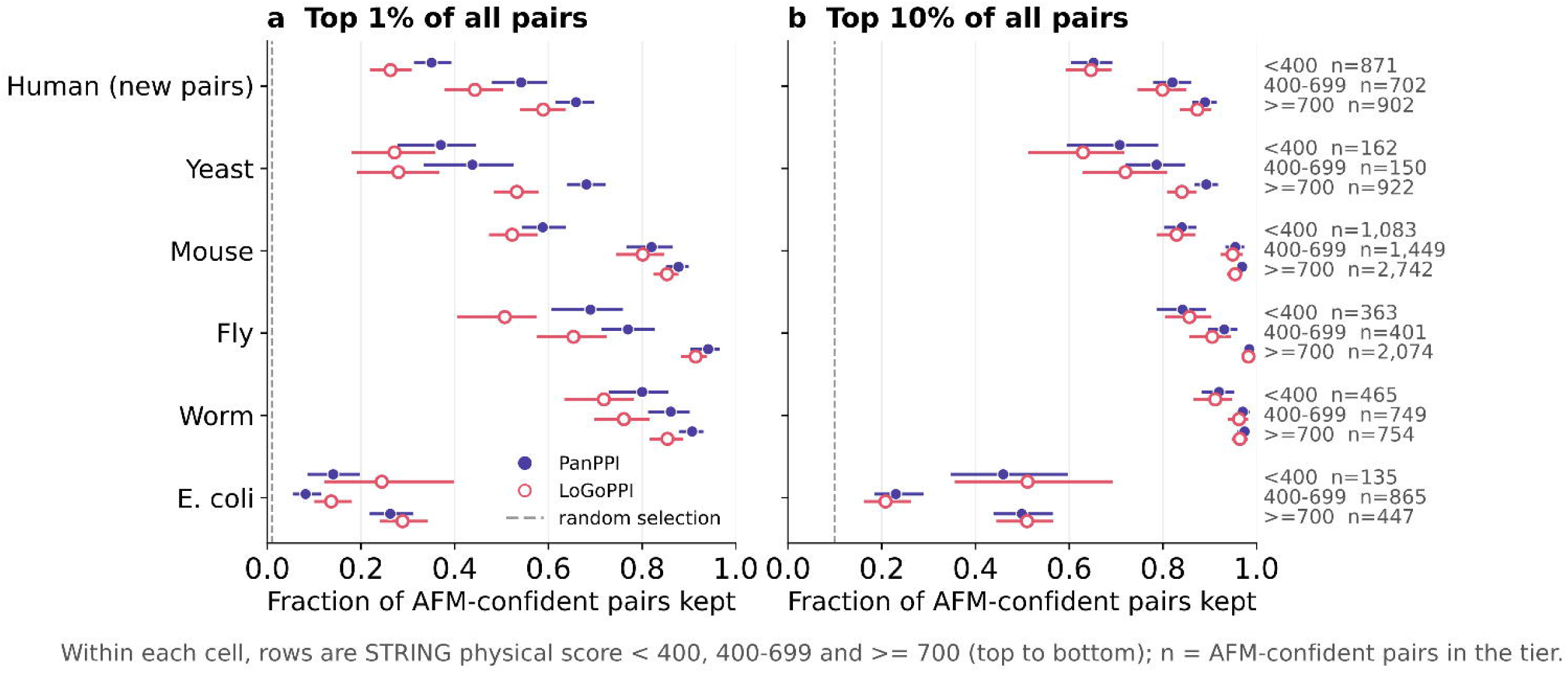

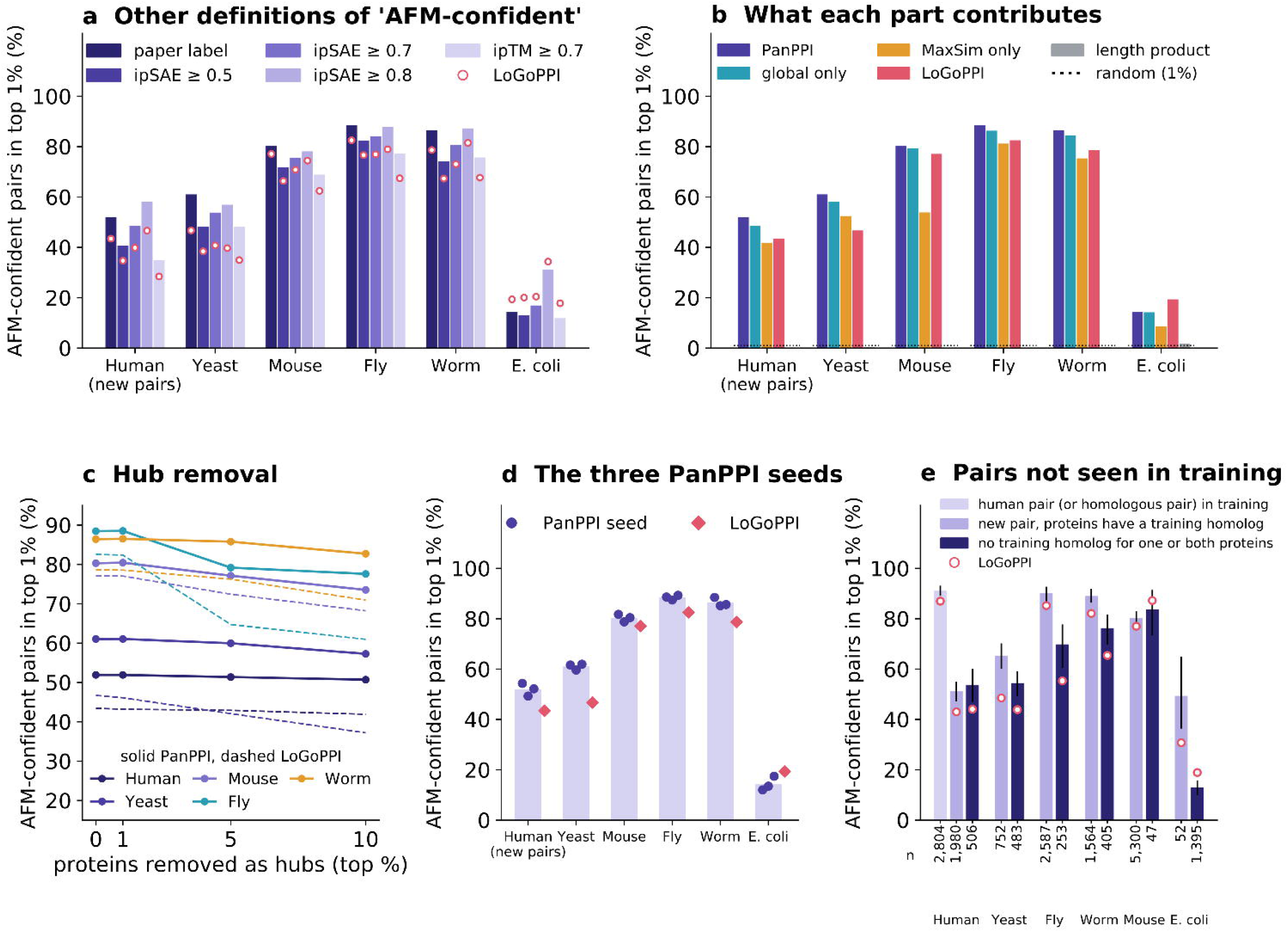

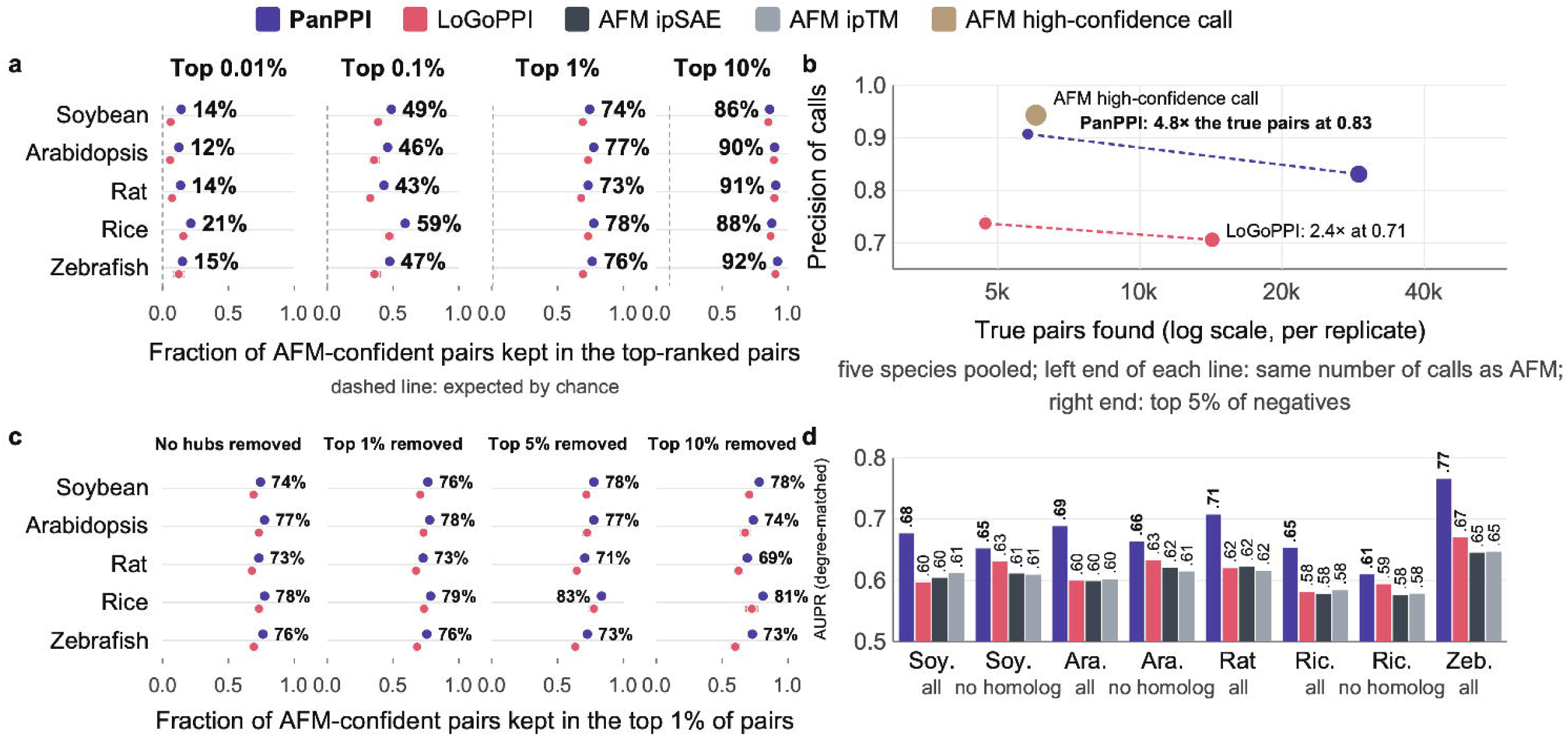

