## Supplementary_Data for "PanPPI: ranking-trained protein embeddings for proteome-scale interaction screening"

### Supplementary Information

**Supplementary Note 1 | Evaluation sets, cell inclusion and *E. coli*.** A cell is one species at one homology level (all proteins, or only proteins with no homolog at 40% identity and 50% query coverage to the human training proteins). Negatives were built to match each protein's positive degree (Havel-Hakimi construction with degree-preserving swaps; three replicates). For the comparison with AlphaFold-Multimer, negatives were low-confidence AlphaFold Database candidates, positives were trimmed to those whose degrees the candidate graph could match, and a cell was used only if its negatives reached 80% of the retained positives and a degree-only score had an AUROC of at most 0.60; this gave 9 cells (5 with all proteins, 4 with no homolog). Trimming kept 38-72% of the STRING positives in these cells (human 72% and 60%, mouse 55% and 39%, fly 45% and 43%, yeast 45% and 38%, worm 45%; all proteins and no homolog, respectively), so the sets favour pairs whose proteins have moderate degree. Precision in Fig. 2c refers to balanced sets: at a prevalence of 0.1%, the threshold operating point (5% false-positive rate) corresponds to a precision of about 0.6% for PanPPI and 0.3% for LoGoPPI (derived from their balanced-set precision of 0.866 and 0.759), which suits a shortlist but not a final call. Trimming enlarges PanPPI's lead over LoGoPPI (mean AUPR difference 0.042 on the untrimmed degree-balanced sets of **Supplementary Fig. 2** against 0.067 on the trimmed sets, over the same nine cells); with all positives kept and the negatives reweighted instead, PanPPI still ranks above LoGoPPI and AlphaFold-Multimer in every species (**Supplementary Table 15**). The degree-only AUROC of every cell is shown in Supplementary Fig. 2d.

In *E. coli*, the AFM-confident pairs have weaker STRING support than in the other species: 31% have a STRING physical score of at least 700, against 75% in yeast, 73% in fly, 52% in mouse and 38% in worm. Retention was lower for both tools even for the *E. coli* pairs with STRING ≥700 (top 1%: PanPPI 26% [22-31], LoGoPPI 29% [24-34], against 66-94% for PanPPI in the eukaryotic sets; Supplementary Fig. 7), so weak STRING support alone does not explain it.

**Supplementary Note 2 | Post-hoc uncertainty and familiarity features add accuracy under species shift, and have limits.** PanPPI can be extended after training with three label-free layers: pair-level GP variance, protein-level familiarity and MaxSim evidence (**Methods**). Added to the calibrated blend cumulatively (B1 to B3), they raise cross-species AUPR on the standard files by 0.020 in yeast [0.017, 0.024], 0.015 in worm and 0.008 in fly, with three seeds and the regression fitted on the other four species; mouse (+0.000 [−0.002, +0.002]) and *E. coli* (+0.001 [−0.006, +0.008]) do not gain, and macro AUPR over the five species rises from 0.868 to 0.877 (**Supplementary Fig. 4a**). The gain does not depend on memorised hubs alone: on the Bernett-style cross-species sets, where each protein is equally frequent in positives and negatives, all five species still gain (B3, +0.004 to +0.019, every interval excluding zero; **Supplementary Fig. 4b**). It does not appear within Bernett's human test split (B3 −0.000 [−0.001, +0.001]; **Supplementary Fig. 4b**; regression fitted on the validation set, the half used for model selection, which is the half with GP variance), and, as a control, a regression fitted on the standard species lowers Bernett AUPR by 0.072 (B1, 0.088**; Supplementary Table 6**). The layers therefore help when a model is applied to a new species, including under hub matching, not when it is applied to held-out proteins of the training species.

The features are informative but they are not an error flag. On the standard files the GP variance predicts misclassifications (AUROC 0.64-0.89), but low score confidence alone predicts them better (0.85-0.94), and variance and confidence are strongly correlated (−0.56 to −0.73). Under the Bernett protocol the relation reverses: variance is higher for confident predictions (correlation +0.38 to +0.53) and predicts errors below chance (AUROC 0.37-0.43; **Supplementary Table 9**), so the regression uses it with the opposite sign. We therefore report these layers as a measured accuracy gain under species shift and do not claim calibrated uncertainty or error detection under leakage control.

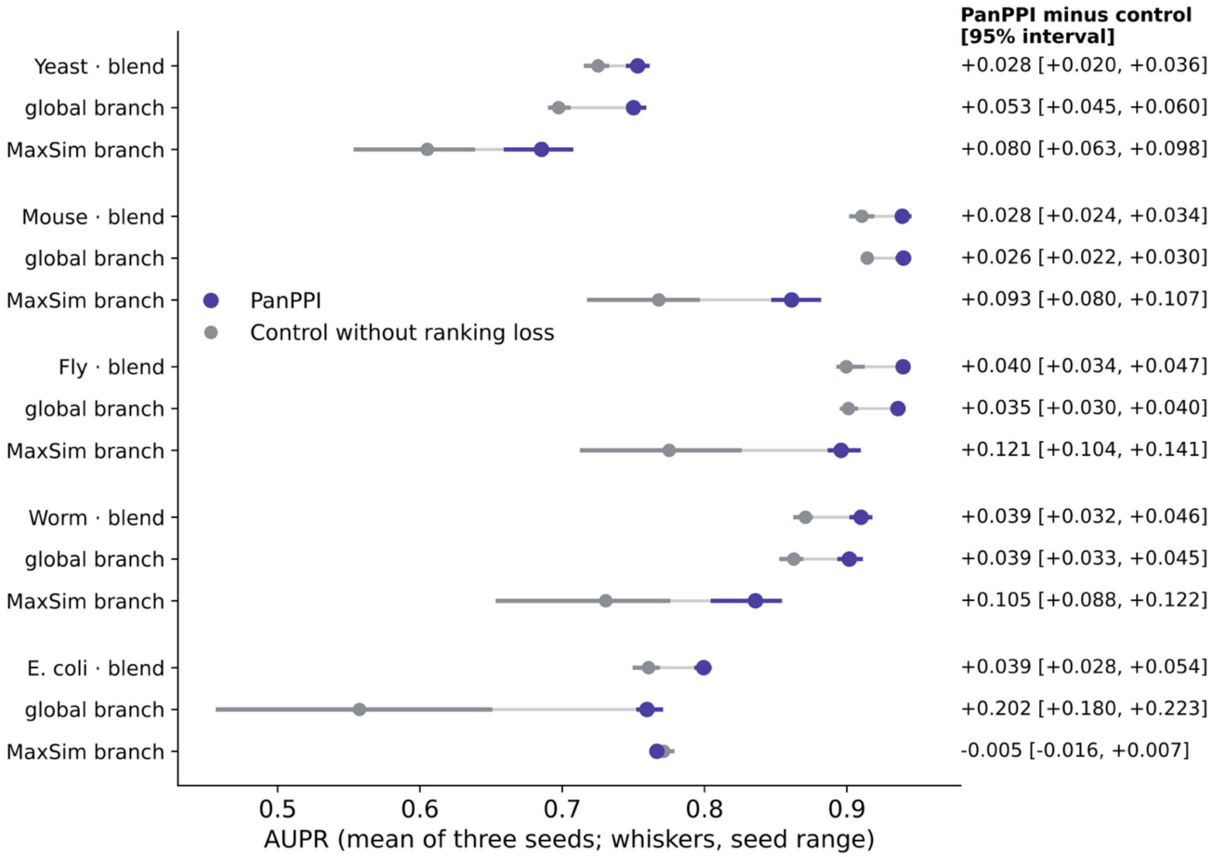


**Supplementary Fig. 1 | A control without the ranking loss.** AUPR of PanPPI and of a control trained with the LoGoPPI recipe in our harness (ranking-loss weight 0) on the five D-SCRIPT test species, for the blend and for each branch. Dots, mean of three seeds; whiskers, seed range; right, PanPPI minus control with 95% paired, protein-clustered bootstrap intervals (1,000 resamples).

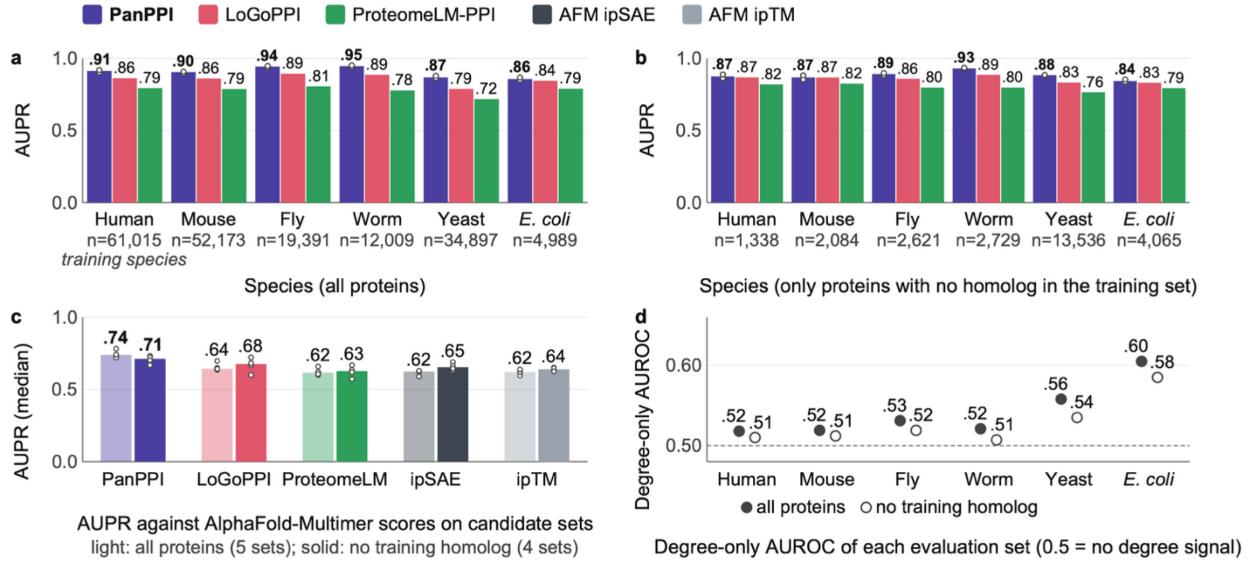


**Supplementary Fig. 2 | Degree-balanced evaluation sets behind Figure 2. a, b**, AUPR of PanPPI (mean of three seeds; circles, individual seeds), LoGoPPI and ProteomeLM-PPI on STRING physical (score ≥700) positives against degree-matched negatives, for all proteins (**a**) and for proteins with no training homolog (**b**); n, positive pairs; human is the training species. **c**, AUPR against AlphaFold-Multimer ipSAE and ipTM on AlphaFold Database candidate sets; light bars, all proteins; solid bars, no training homolog; bars, medians over sets; circles, single sets. STRING and PDB labels are of the type PanPPI and LoGoPPI were trained on; AlphaFold-Multimer was not (**Fig. 2d** for verified contacts). **d**, AUROC of a degree-only score for every set in **a** and **b**; 0.5 means no degree signal.

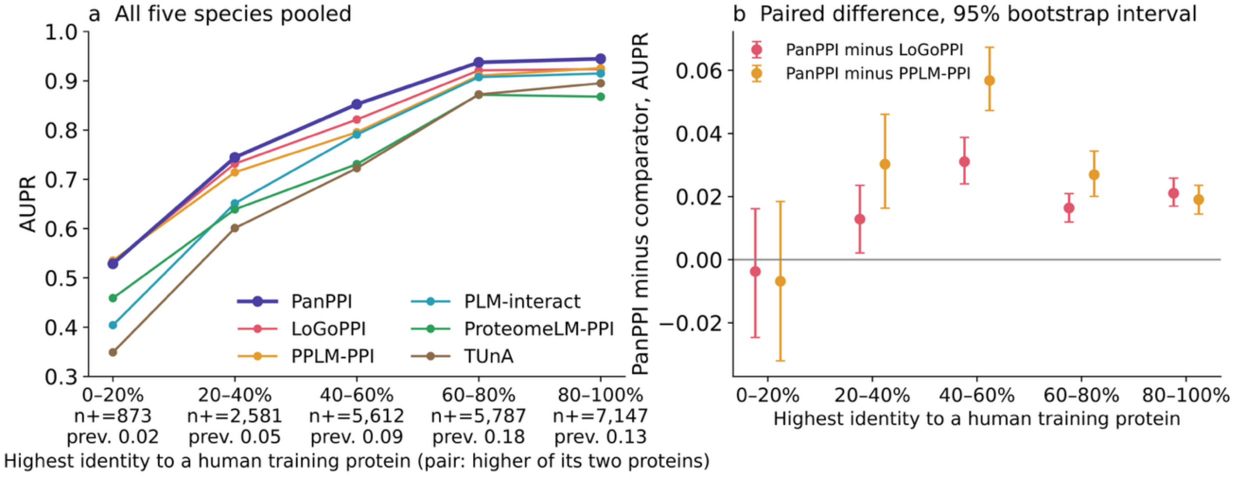


**Supplementary Fig. 3 | Accuracy by similarity to the human training proteins.** Pairs from the five D-SCRIPT test species, binned by the highest sequence identity of a pair's more similar protein to any human training protein (MMseqs2; no hit counts as 0). **a**, AUPR of each tool, pooled over the five species; n+, positive pairs; prev., prevalence in the bin, on which AUPR depends. **b**, PanPPI minus LoGoPPI and minus PPLM-PPI with 95% paired, protein-clustered bootstrap intervals (1,000 resamples). In the 0-20% bin the three tools do not differ; the margin is largest at 40-60%.

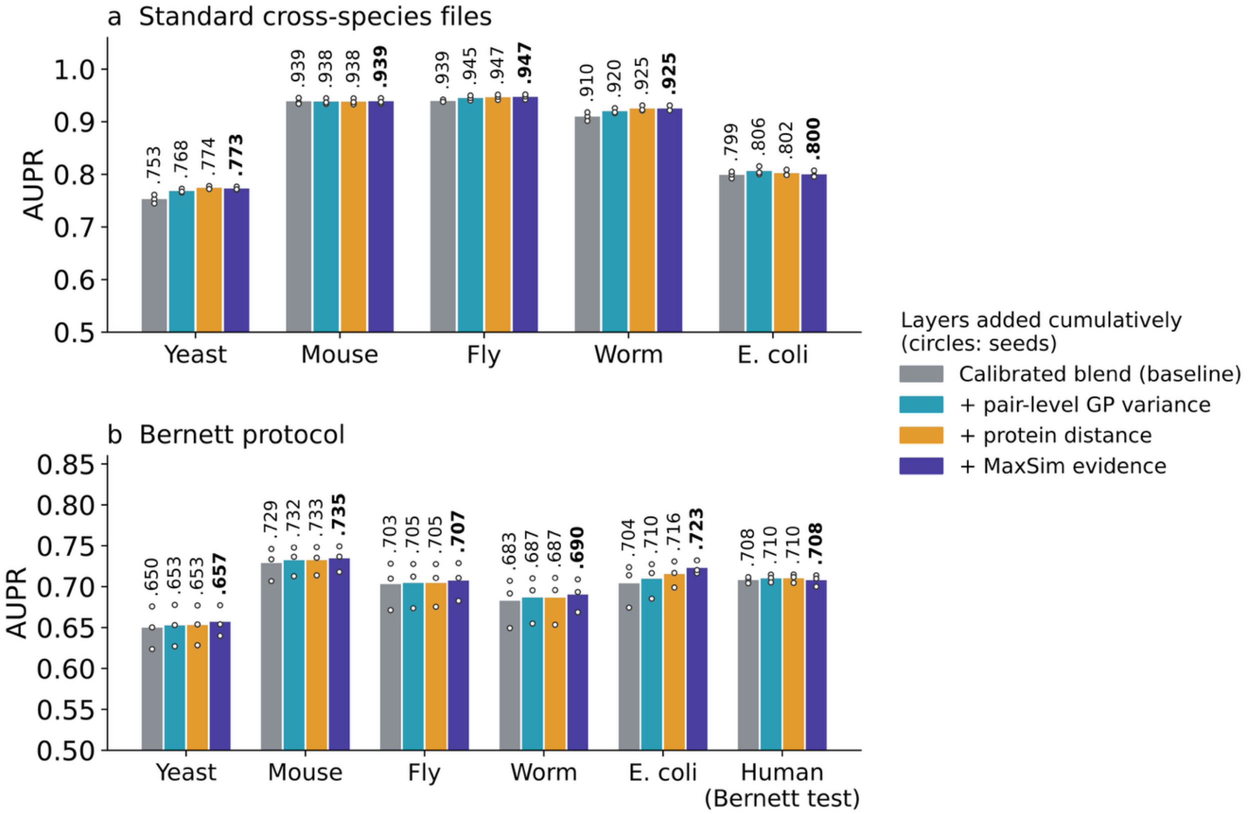


**Supplementary Fig. 4 | Effect of the uncertainty and familiarity features on AUPR.** AUPR of the calibrated blend (grey, baseline) and of the blend with three layers added cumulatively: pair-level GP variance (teal), protein distance to the nearest training proteins (orange) and MaxSim evidence (purple). Bars, mean of three seeds; circles, seeds; axes start at 0.5. **a**, Standard cross-species files, regression fitted on the other four species. **b**, Bernett protocol: Bernett-style cross-species sets (regression fitted on the other four) and the human Bernett test (regression fitted on the Bernett validation set). A regression fitted on the standard species and applied to the Bernett test is a control; it and the paired intervals are in Supplementary Table 6.

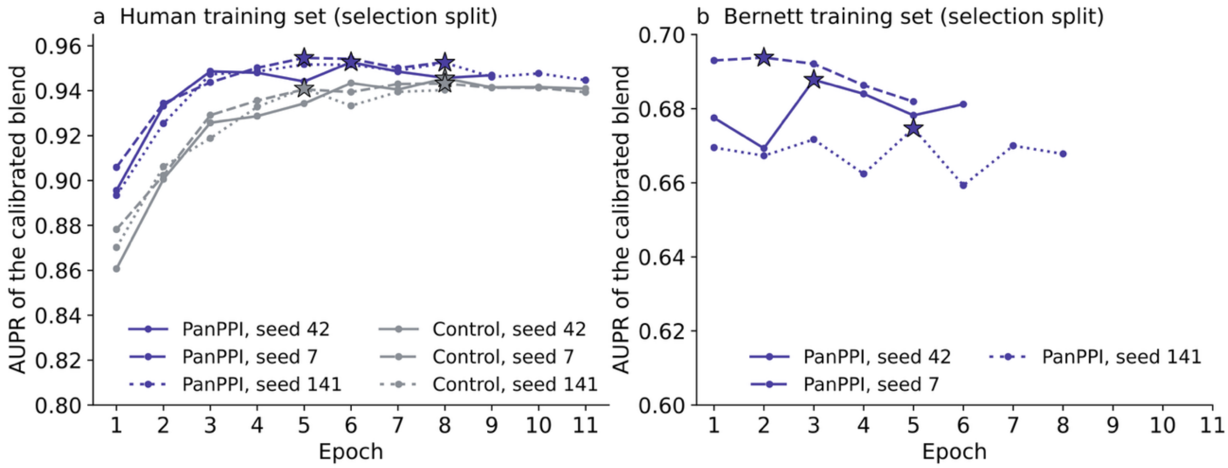


**Supplementary Fig. 5 | Training curves and early stopping.** AUPR of the calibrated blend on the selection split after each epoch. **a**, Human training data: PanPPI (purple) and the control without the ranking loss (grey), three seeds each (line styles). **b**, PanPPI trained on the Bernett split, three seeds. Stars mark the epoch kept; each run stopped after three epochs without improvement and none reached the 14-epoch ceiling.

**Supplementary Note 3 | Models trained on the Bernett split and verified contacts.** The Bernett-trained PanPPI and LoGoPPI were scored on the 30,024 human candidate pairs from which Fig. 2d is drawn (**Supplementary Table 7**); the contact analysis uses 3,595 positives and 15,852 negatives of them. Training on the leakage-controlled split did not close the gap to AlphaFold-Multimer on direct contacts: for contact pairs new to the human-trained models, PanPPI-Bernett reached an AUPR of 0.639 [0.616, 0.670] against 0.644 for the human-trained PanPPI and 0.796 for AFM ipSAE. For pairs seen by the human-trained models, the Bernett-trained models scored lower (0.700 against 0.895 for PanPPI), as expected if the human-trained models partly memorise pairs. For same-entry pairs without contact, PanPPI-Bernett (0.634) remained above AFM ipSAE (0.588). The seen and new labels refer to the human-trained training set; overlap with the Bernett training set was not recomputed.

**Supplementary Note 4 | GO- and Pfam-matched hard negatives.** We repeated LoGoPPI's hard-negative evaluation (its **Supplementary Fig. 4**) on the Bernett test set with the Bernett-trained models. A negative pair is hard if its two proteins share a direct, positively asserted GO cellular-component annotation (GOA human, aspect C; excluding the root, terms of the subset gocheck_do_not_annotate and protein-containing complex with its is_a descendants; go-basic data version 2026-07-26) or an identical Pfam accession (UniProt release 2026_04). Positives are drawn from the positives that also share, with replacement, to the number of hard negatives; hard negatives and size-matched random negatives are resampled with replacement (2,000 replicates; the same positives in both comparisons), and PanPPI is the mean of its three seeds' own AUPR. Our counts reproduce LoGoPPI's: 19,490 positive and 13,705 negative pairs share a cellular component (LoGoPPI: 19,518 and 13,714) and 1,483 and 286 share a Pfam domain (identical), and LoGoPPI-Bernett reaches AUPR 0.644 and 0.645 on the two hard sets (reported: about 0.64). All models lose accuracy on hard negatives. With shared cellular components PanPPI is ahead of all three comparators (AUPR 0.677 against 0.644 for LoGoPPI, 0.658 for PLM-interact and 0.660 for TUnA; **Supplementary Fig. 6** and **Supplementary Table 10**). With shared Pfam domains it is level with LoGoPPI (0.645 against 0.645) and ahead of PLM-interact (0.595) and TUnA (0.547), with wide intervals because only 286 negatives qualify.

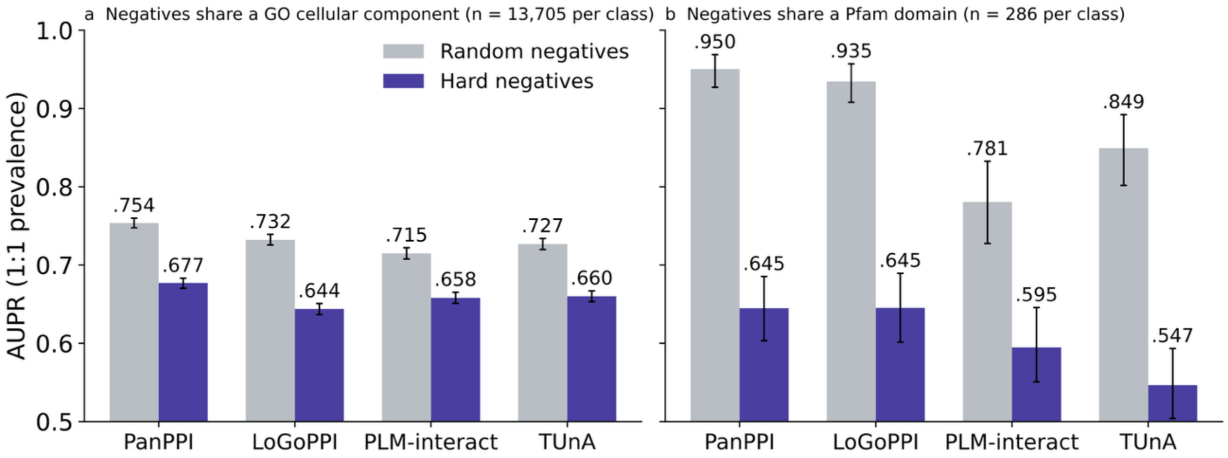


**Supplementary Fig. 6 | GO- and Pfam-matched hard negatives.** AUPR of the Bernett-trained models on the Bernett test set with size-matched random negatives (grey) and hard negatives (purple), with positives drawn from positives that also share the annotation (1:1 prevalence); PanPPI, mean of three seeds; whiskers, 2.5-97.5 percentiles of 2,000 bootstrap replicates. Hard negatives share a GO cellular component (**a**) or a Pfam domain (**b**).

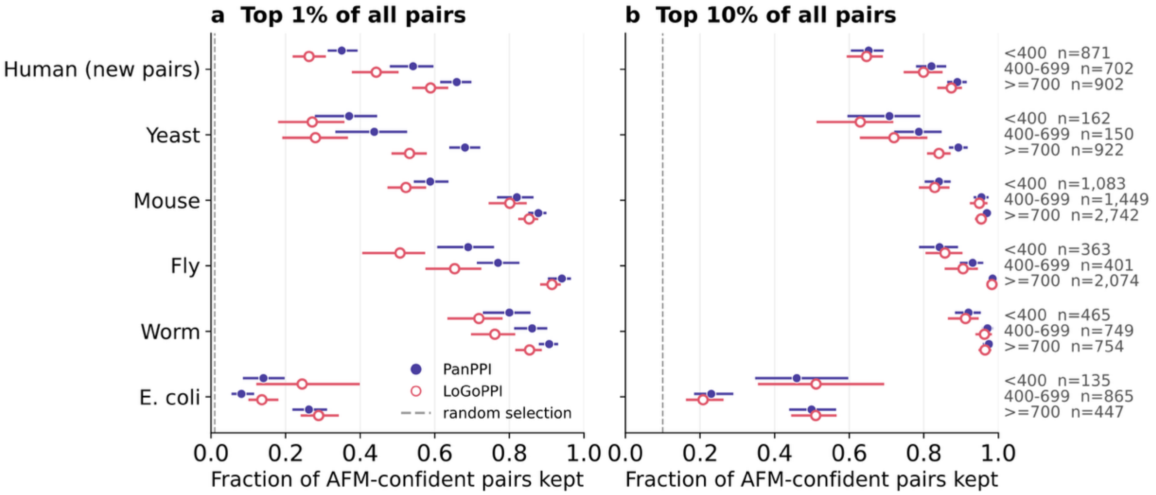


Within each cell, rows are STRING physical score < 400, 400-699 and >= 700 (top to bottom); n = AFM-confident pairs in the tier.

**Supplementary Fig. 7 | Retention of AFM-confident pairs by STRING support.** Share of AFM-confident pairs kept in the top 1% (**a**) and top 10% (**b**) of all pairs, split by the pair's STRING physical score (below 400, 400-699, 700 or more; top to bottom within each cell). PanPPI (mean of three seeds, filled) and LoGoPPI (open) with 95% protein-clustered bootstrap intervals (300 resamples); the dashed line is the share expected from random selection; n is the number of AFM-confident pairs in each tier. Retention rises with STRING support, so part of it reflects agreement with STRING, but in the five eukaryotes 35-80% of the pairs with a STRING score below 400 are in the top 1% (**Supplementary Table 16**). Human: pairs new to the model.

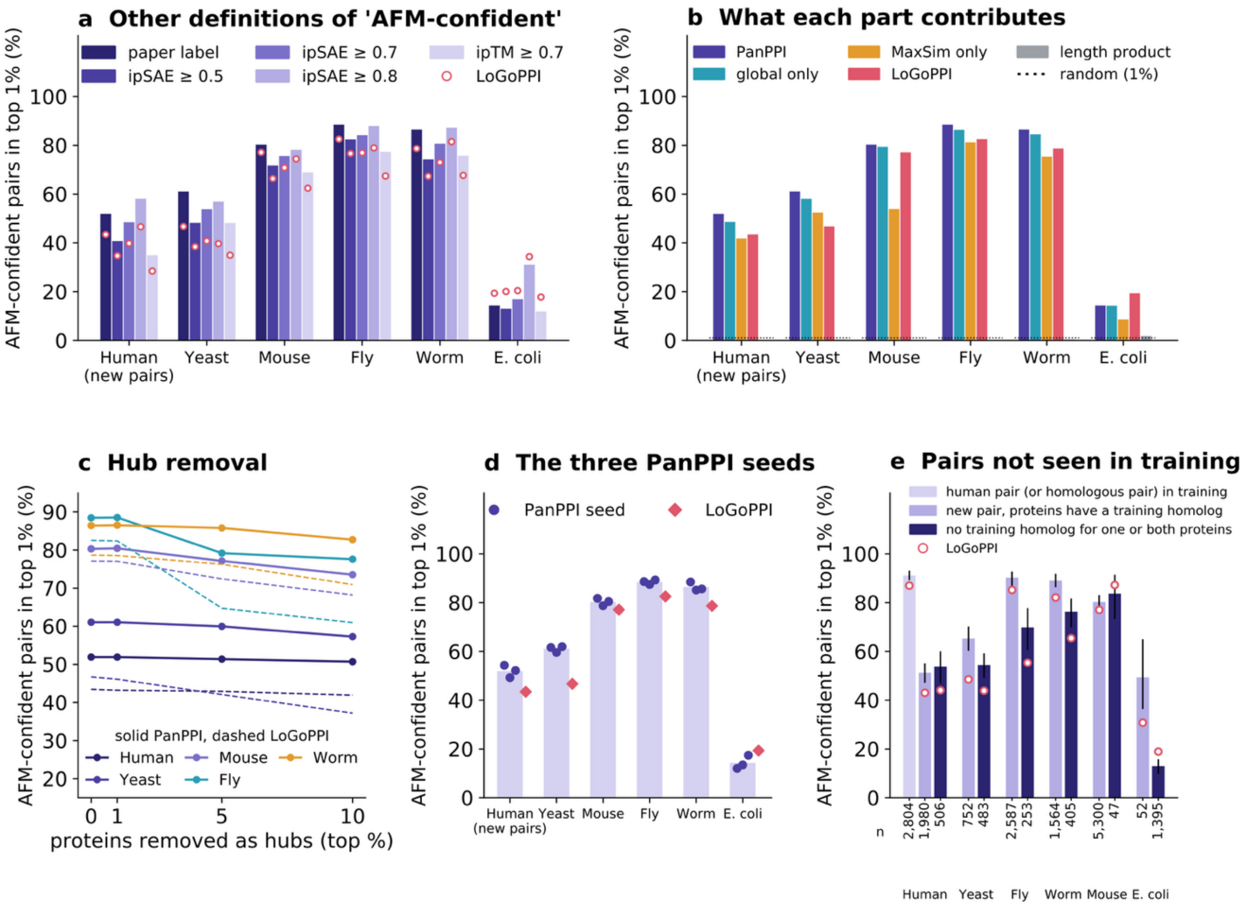


**Supplementary Fig. 8 | Retention of AFM-confident pairs: other definitions, the parts of PanPPI, hub removal, seeds and unseen pairs.** All panels show the share of AFM-confident pairs among the top 1% of all pairs of the proteome (human: pairs new to the model, except in **e**). **a**, Retention when 'AFM-confident' is defined by ipSAE (≥0.5, ≥0.7 or ≥0.8) or ipTM (≥0.7) alone, against the label used in the paper (ipSAE ≥0.6 and pDockQ2 ≥0.23); PanPPI, bars; LoGoPPI, circles. **b**, PanPPI, its global and MaxSim branches alone, LoGoPPI and a protein-length baseline; dotted line, random selection. **c**, Retention after removing pairs that involve the top 1%, 5% or 10% of proteins by number of STRING-linked candidate partners (solid, PanPPI; dashed, LoGoPPI). **d**, The three PanPPI seeds (circles) and their mean (bar); diamonds, LoGoPPI. **e**, Pairs not seen in training. Human pairs are split by overlap with the training set (pair or homologous pair in training; new pair whose proteins both have a training homolog; new pair with one protein without a homolog; the nine pairs with neither protein homologous are not shown); other species are split by whether either protein has a homolog among the human training proteins (≥40% identity, ≥50% query coverage). Whiskers, 95% protein-clustered bootstrap intervals (200 resamples); numbers under the bars, AFM-confident pairs; LoGoPPI as in **a**.

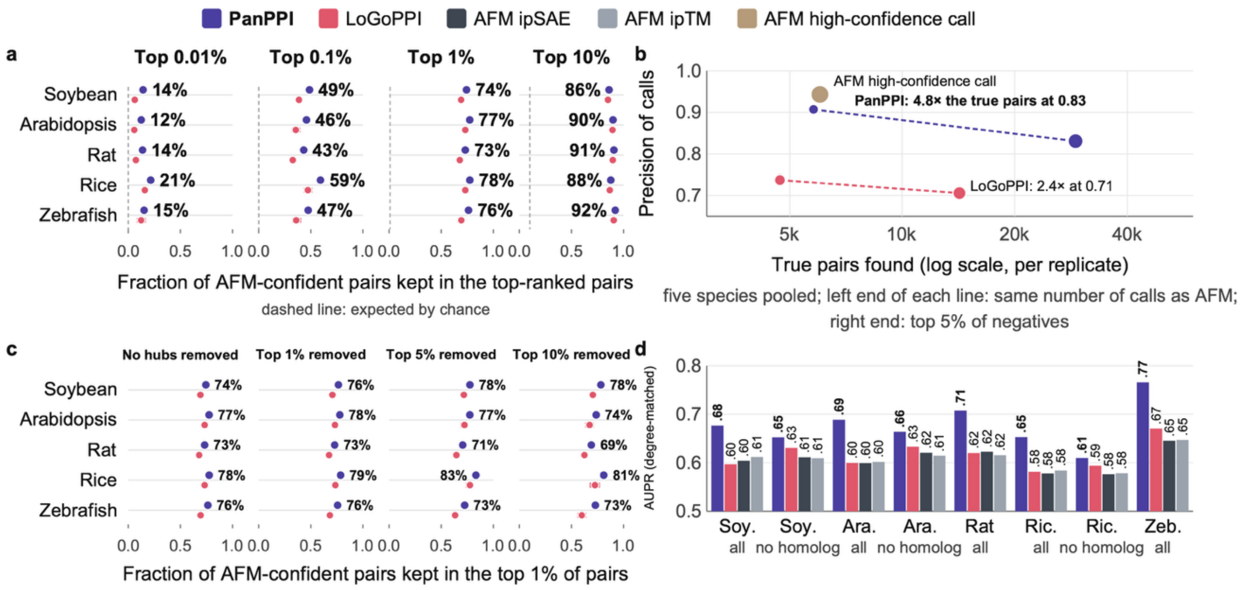


**Supplementary Fig. 9 | Five further species: soybean, *Arabidopsis*, rat, rice and zebrafish.** The analyses of **Fig. 2** for the five proteomes of the AlphaFold Database complexes resource with the most AFM-confident pairs after the main species, a rule fixed before any model was scored (not part of the main figure), drawn as in **Fig. 2** (PanPPI, mean of three seeds; LoGoPPI, released model); AFM-confident pairs from the AlphaFold Database complexes resource, STRING v12 as for the main species. **a**, Share of AFM-confident pairs (4,219-14,100 per species) among the top 0.01%, 0.1%, 1% and 10% of all pairs; dashed line, expected by chance; whiskers, 95% protein-clustered bootstrap intervals (300 resamples). **b**, Precision against STRING physical pairs of the AFM high-confidence call, of PanPPI and LoGoPPI calling as many pairs as AFM (left end of each line) and of each tool above the 95th percentile of its negatives (right end); five species and three negative replicates pooled; true pairs per replicate on a log axis. **c**, As **a** for the top 1%, after removing pairs that involve the top 1%, 5% or 10% of proteins by number of STRING-linked candidate partners. **d**, AUPR on degree-matched sets of STRING physical pairs against low-confidence candidate negatives, for all proteins and for proteins with no homolog among the human training proteins; a set is shown only if its negatives reach 80% of the retained positives and a degree-only score has an AUROC of at most 0.60, which excludes the no-homolog sets of rat and zebrafish.
